# A Neural Basis for Mutant ATAXIN-1 Induced Respiratory Dysfunction in Mouse Models of Spinocerebellar Ataxia Type 1

**DOI:** 10.1101/2024.08.20.608114

**Authors:** Alyssa Soles, Jessica Grittner, Kaia Douglas, Praseuth Yang, Ryan Barnett, Christine Chau, Roj Cosiquien, Lisa Duvick, Orion Rainwater, Shannah Serres, Harry Orr, Brendan Dougherty, Marija Cvetanovic

## Abstract

Spinocerebellar ataxia type 1 (SCA1), a dominantly inherited neurodegenerative disorder caused by an expanded trinucleotide repeat in the *ATAXIN-1 (ATXN1)* gene, is characterized by motor dysfunction, cognitive impairment, and death from compromised swallowing and respiration. To delineate specific cell types that contribute to respiratory dysfunction, we utilized the floxed conditional knock-in *f-ATXN1^146Q/2Q^* mouse. Whole body plethysmography during spontaneous respiration and respiratory challenge showed that *f-ATXN1^146Q/2Q^* mice exhibit a spontaneous respiratory phenotype characterized by elevated respiratory frequency, volumes, and respiratory output. Consequently, the ability of *f-ATXN1^146Q/2Q^* mice to increase ventilation during the challenge is impaired. To investigate the role of mutant *ATXN1* expression in neural and skeletal muscle lineages, *f-ATXN1^146Q/2Q^* mice were bred to *Nestin-Cre* and *Acta1-Cre* mice respectively. These analyses revealed that the abnormal spontaneous respiration in *f-ATXN1^146Q/2Q^* mice involved two aspects: a behavioral phenotype in which SCA1 mice exhibit increased motor activity during respiratory testing and functional dysregulation of central respiratory control centers. Both aspects of spontaneous respiration were partially ameliorated by removing mutant *ATXN1* from neural, but not skeletal muscle, cell lineages.

## Introduction

Breathing is an essential behavior that is based on complex neural circuitry. Respiratory dysfunctions are commonly found in patients with neurodegenerative disease, and respiratory complications are often linked to their cause of death (Bloem et al., 2021; Gaig & Iranzo, 2012; Hobson & Meara, 2018; Huebra et al., 2019; Larson et al., 2023; Sørensen & Fenger, 1992). Examining the neural basis of respiratory dysfunction associated with disease affords a unique opportunity to further understand how the nervous system regulates breathing and how nervous system dysfunction impacts this vital function.

Spinocerebellar ataxia type 1 (SCA1) is a dominantly inherited adult-onset neurodegenerative disease caused by an expanded trinucleotide repeat within the *ATAXIN-1* (*ATXN1*) gene. SCA1 is characterized by motor dysfunction, cognitive impairment, and premature death from compromised breathing and swallowing (Diallo et al., 2018; Genis et al., 1995; Robitaille et al., 1995; Sasaki et al., 1996). In a recent study, Coarelli et al. reported dysfunction of the larynx and diaphragm muscles that is associated with upper motor neuron pathology in a cohort of SCA1 patients with respiratory related complications (Coarelli et al., 2023). Whether motor neuron pathology is a cause or consequence of larynx and diaphragm muscle dysfunction in SCA1 is unknown.

Mouse models of SCA1 are a valuable tool for understanding the underlying biology of how abnormal polyglutamine expansions in *ATXN1* perturb physiology. The *Atxn1^154Q/2Q^* SCA1 mouse model exhibits a progressive dysfunction of spontaneous respiration with associated pathology similar to SCA1 patients including motor neuron pathology and gliosis in the brainstem and spinal cord (Orengo et al., 2018). In addition, respiratory dysfunction in a mouse model of spinocerebellar ataxia type 7 (SCA7) is associated with neurodegeneration and inflammation in phrenic and hypoglossal motor neurons of the spinal cord and brainstem respectively (Fusco et al., 2021).

Recently, a humanized conditional SCA1 knock-in mouse model was developed, the *f-ATXN1^146Q/2Q^* line, which expresses the human coding region of expanded *ATXN1* flanked with lox sites in place of the endogenous mouse *Atxn1* gene (Duvick et al., 2024). This model allows for the contribution of specific regions and cell types to disease phenotypes to be examined via deletion of *ATXN1* with Cre-recombinase. While SCA1 is traditionally thought of as a neurodegenerative disease, characterization of this model indicates abnormal muscle function including a reduction in force likely caused by neuronal dysfunction earlier in disease and later by an intrinsic weakening of muscle fibers. Automatic respiration is largely regulated by a collection of nuclei in the brainstem that form a central pattern generator to pattern muscle activity. As the integrity of the nervous system and skeletal muscle are all compromised in mouse models of SCA1, in this study, we utilized the *f-ATXN1^146Q/2Q^* SCA1 mouse to understand the extent to which expression of mutant ATXN1 in these regions contributes to respiratory dysfunction.

## Results

### *f-ATXN1^146Q/2Q^* mice exhibit a progressive dysfunction of spontaneous and challenged respiration

Respiratory function was assessed in unanesthetized and unrestrained *f-ATXN1^146Q/2Q^* and *Atxn1^2Q/2Q^* (wild-type) mice using whole body barometric plethysmography (WBP). Breathing was recorded during normoxia (21% O_2_, balance N_2_) to examine spontaneous respiration and during a combined hypoxic and hypercapnic condition (10% O_2_, 5% CO_2_, balance N_2_) to examine the ability of mice to regulate respiration in response to challenge. Figure 1A depicts representative WBP flow traces from two-hour recordings of spontaneous respiration followed by a five-minute hypoxic and hypercapnic challenge at 35 weeks in *f-ATXN1^146Q/2Q^* and wild-type (WT). Dashed line a marks the beginning of the last 30 minutes of spontaneous respiration and b marks the beginning of the challenge. By 35 weeks *f-ATXN1^146Q/2Q^* mice exhibited a remarkably consistent respiratory pattern characterized by a high frequency and amplitude of respiratory flow traces during spontaneous respiration without the typical respiratory waveform variability observed in WT animals (Figure 1A). All respiratory parameters calculated in figures 1B-1D were assessed from a three-minute segment of respiration within the last 30 minutes of spontaneous respiration. This segment was selected by examining the entire final 30 minutes of spontaneous respiration for a representative segment that depicted the most frequent and consistent respiratory pattern in individual mice. Hypoxic and hypercapnic challenge was analyzed from the final one-minute of the challenge across all animals. Respiratory variables including respiratory rate, inspiratory time, expiratory time, tidal volume, and minute ventilation were automatically calculated with Ponemah (DSI, St. Paul, MN). Central drive, or tidal volume production during the inspiratory period, was manually compared as a proxy for neural drive to respiratory muscles (Georgopoulos et al., 2024; Morgan et al., 2014).

**Figure 1:**
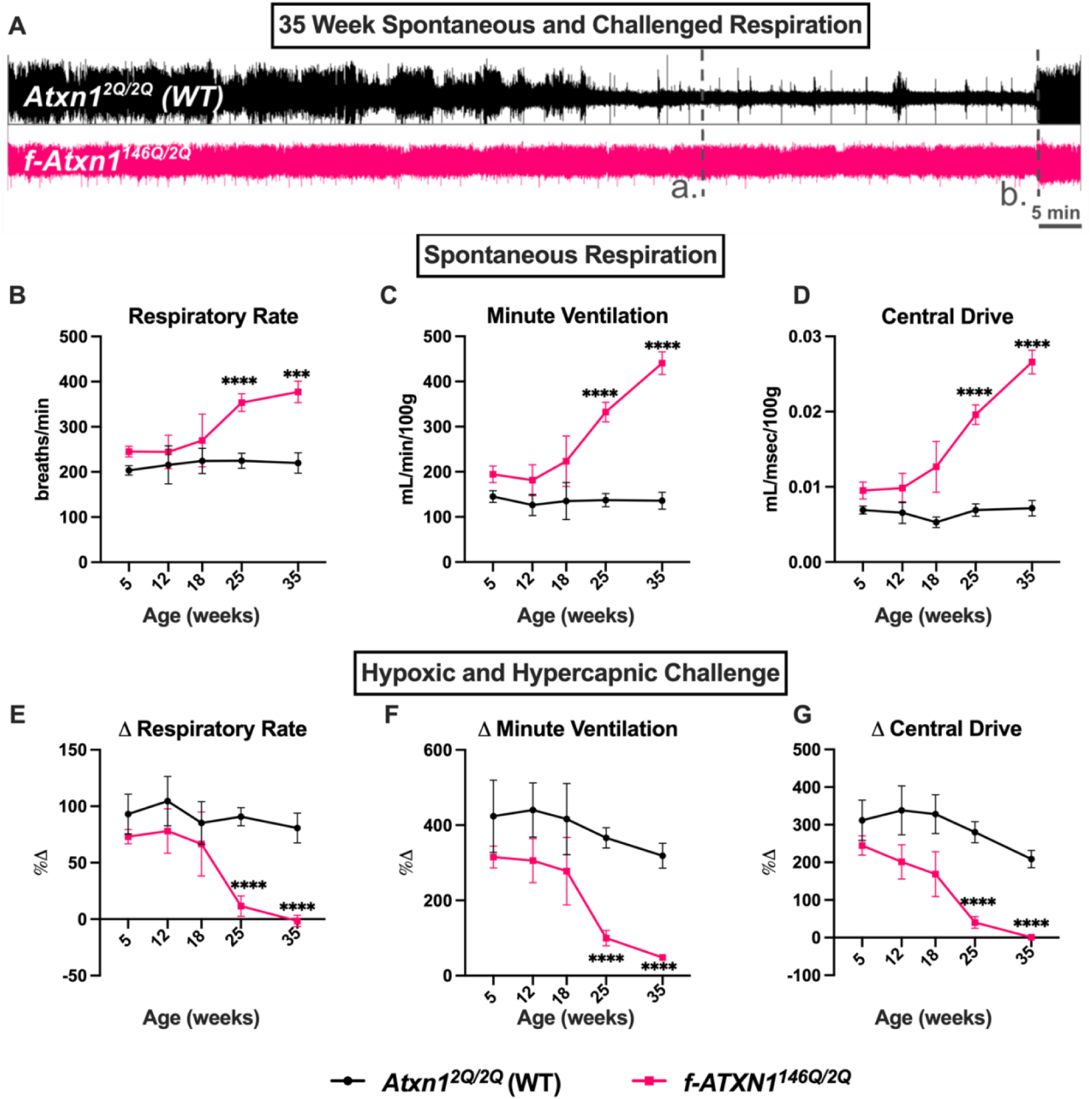
*f-ATXN1^146Q/2Q^* mice exhibit a progressive dysfunction of spontaneous and challenged respiration. (**A**) Whole body plethysmography flow traces demonstrating representative two-hour recordings of spontaneous respiration followed by a five minute hypoxic and hypercapnic challenge in 35 week WT and *f-ATXN1^146Q/2Q^* mice. The dashed line labeled with a. marks the start of the last 30 minutes of spontaneous respiration while b. marks the beginning of the challenge. Spontaneous respiration was quantified in the segment between a. and b. as respiratory rate (**B**), minute ventilation (**C**) and central drive (**D**). The response of mice to the final minute of the hypoxic and hypercapnic challenge was assessed as the percentage change in respiratory rate (**E**), minute ventilation (**F**) and central drive (**G**) from spontaneous levels. Statistics for B-G were assessed with a repeated measures mixed effects analysis with Tukey’s and Geisser-Greenhouse corrections. *\*\*\** p<0.001 **** p<0.0001. Sample sizes at each age for WT are 5wk=8, 12wk=6, 18wk=5, 25wk=24, 35wk=21 and for *f-ATXN1^146Q/2Q^* are 5wk=10, 12wk=7, 18wk=5, 25wk=19, 35wk=12.

During the early stages of disease, at 5 and 12 weeks of age, there were no significant differences in spontaneous or challenged respiration between *f-ATXN1^146Q/2Q^* and WT mice (Figure 1, Supplementary figure 1). At 18 weeks, a nonsignificant trend emerged in which *f-ATXN1^146Q/2Q^* mice displayed an elevated respiratory rate, minute ventilation, and central drive. By 25 weeks, *f-ATXN1^146Q/2Q^* mice diverged from WT animals and exhibited a significantly elevated frequency, volume, and central drive during spontaneous respiration (Figure 1B-C, Supplementary figure 1). Consequently, *f-ATXN1^146Q/2Q^* mice failed to significantly increase the central drive of respiration from spontaneous levels during the hypoxic and hypercapnic challenge (Figure 1E-F, Supplementary figure 1).

*F-ATXN1^146Q/2Q^* mice exhibit a failure to gain body weight that manifests as a 50% reduction in body weight by 35 weeks (Supplemental figure 2A). Given the potential role of mouse *Atxn1* in the development of lung tissue (Lee et al. 2011), we dissected and measured the weight of lungs from *f-ATXN1^146Q/2Q^* and WT animals. At 35 weeks, *f-ATXN1^146Q/2Q^* mice exhibit a reduction in lung mass by about 25% compared to WT littermates (Supplemental figure 2B). Histological analysis with hematoxylin and eosin of lungs extracted at 40 weeks from WT and *f-ATXN1^146Q/2Q^* mice did not show signs of fibrosis or inflammation that would contribute to abnormal respiration (data not shown).

### Spontaneous respiration in aged *f-ATXN1^146Q/2Q^* mice is similar to respiration challenged with hypoxia and hypercapnia

We next examined the effect of aging on the respiratory output of *f-ATXN1^146Q/2Q^* and WT mice during spontaneous respiration and respiration challenged simultaneously with hypoxia and hypercapnia. We compared central drive, a proxy for the neural drive to respiratory muscles, at the earliest and the latest age assessed. At 5 weeks, both *f-ATXN1^146Q/2Q^* and WT mice are able to increase respiratory output in response to the hypoxic and hypercapnic challenge (Figure 2). At 35 weeks, *f-ATXN1^146Q/2Q^* mice exhibit a largely increased central drive during spontaneous respiration that is essentially equal to their respiratory output during the hypoxic and hypercapnic challenge (Figure 2A). Consequently, *f-ATXN1^146Q/2Q^* mice fail to increase central drive during the challenge at 35 weeks (Figure 2B).

**Figure 2:**
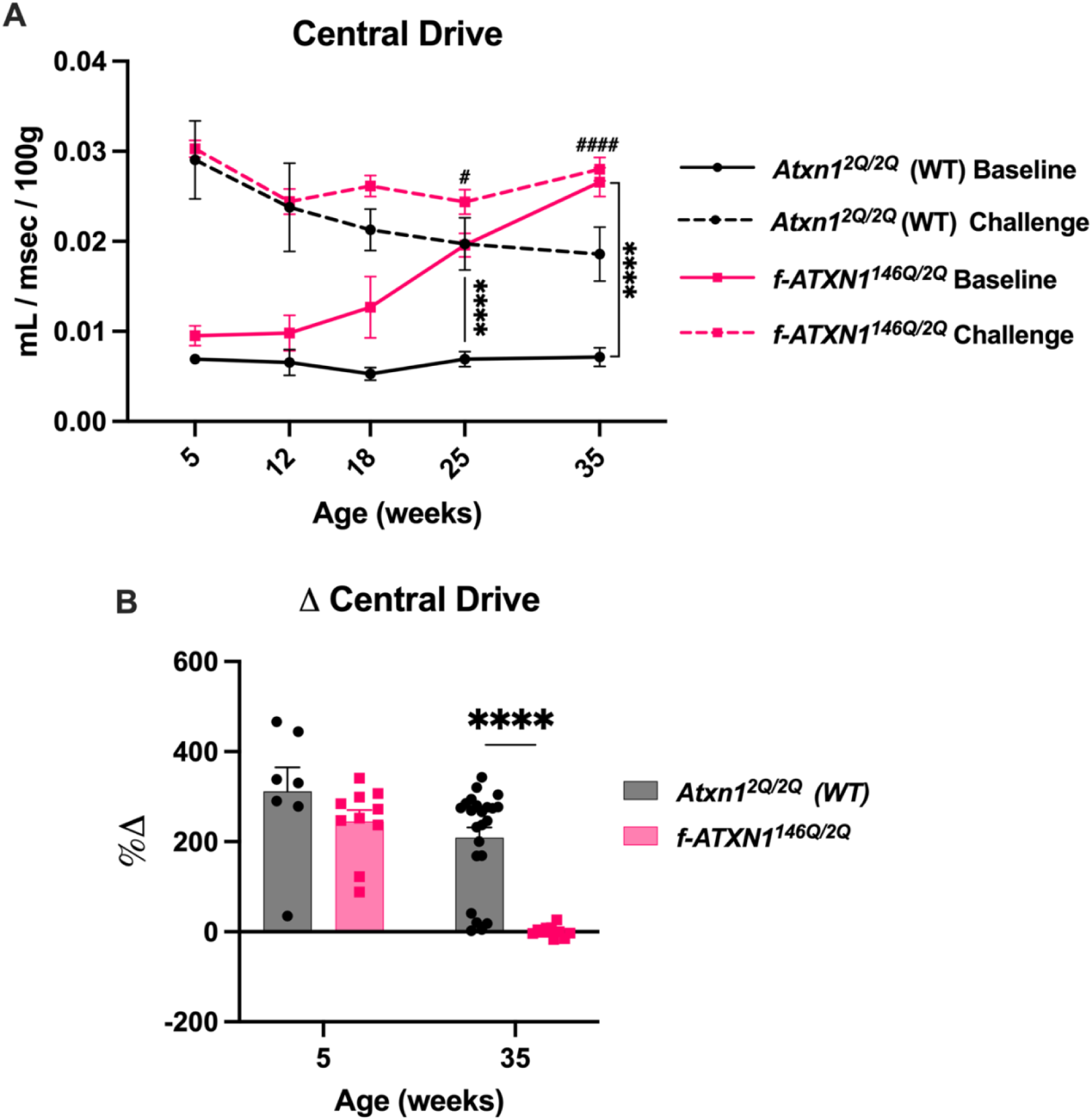
Spontaneous respiration in aged *f-ATXN1^146Q/2Q^* mice is similar to respiration challenged with hypoxia and hypercapnia. (**A**) Central drive during spontaneous respiration and challenged respiration across all ages assessed was plotted in WT and *f-ATXN1^146Q/2Q^* mice. (**B**) Central drive percentage change from baseline levels during the respiratory challenge. Repeated measures mixed effects analysis with Tukey’s and Geisser-Greenhouse corrections. Asterisks (*) denotes significance between groups for spontaneous central drive while number signs (#) denote significance between groups for challenged central drive. # p<0.05, ****/#### p<0.0001. Sample sizes at each age for WT are 5wk=8, 12wk=6, 18wk=5, 25wk=24, 35wk=21 and for *f-ATXN1^146Q/2Q^* are 5wk=10, 12wk=7, 18wk=5, 25wk=19, 35wk=12.

### Increased motor activity during WBP is ameliorated with isoflurane sedation indicating functional dysregulation of respiration in *f-ATXN1^146Q/2Q^* mice

Given the striking increase in the amplitude and frequency of respiratory traces and the quantification of respiratory parameters of 35-week-old *f-ATXN1^146Q/2Q^* mice during spontaneous respiration (Figure 1A), we assessed animal behavior during the acquisition of spontaneous respiration with WBP at 35 weeks. Videography from WBP acquisition was analyzed for movement in the first 30 minutes and last 30 minutes of spontaneous WBP acquisition. During the first 30 minutes of spontaneous respiration, WT and *f-ATXN1^146Q/2Q^* mice exhibited a comparable movement duration (Figure 3A). In the final 30 minutes of spontaneous WBP acquisition, WT animals demonstrated a reduction in movement duration while *f-ATXN1^146Q/2Q^* mice failed to reduce motor activity and exhibited levels of movement similar to the first 30 minutes of WBP acquisition (Figure 3A). This data indicates the behavioral dysregulation during WBP acquisition is an aspect of respiratory dysfunction in 35-week-old *f-ATXN1^146Q/2Q^* mice.

**Figure 3:**
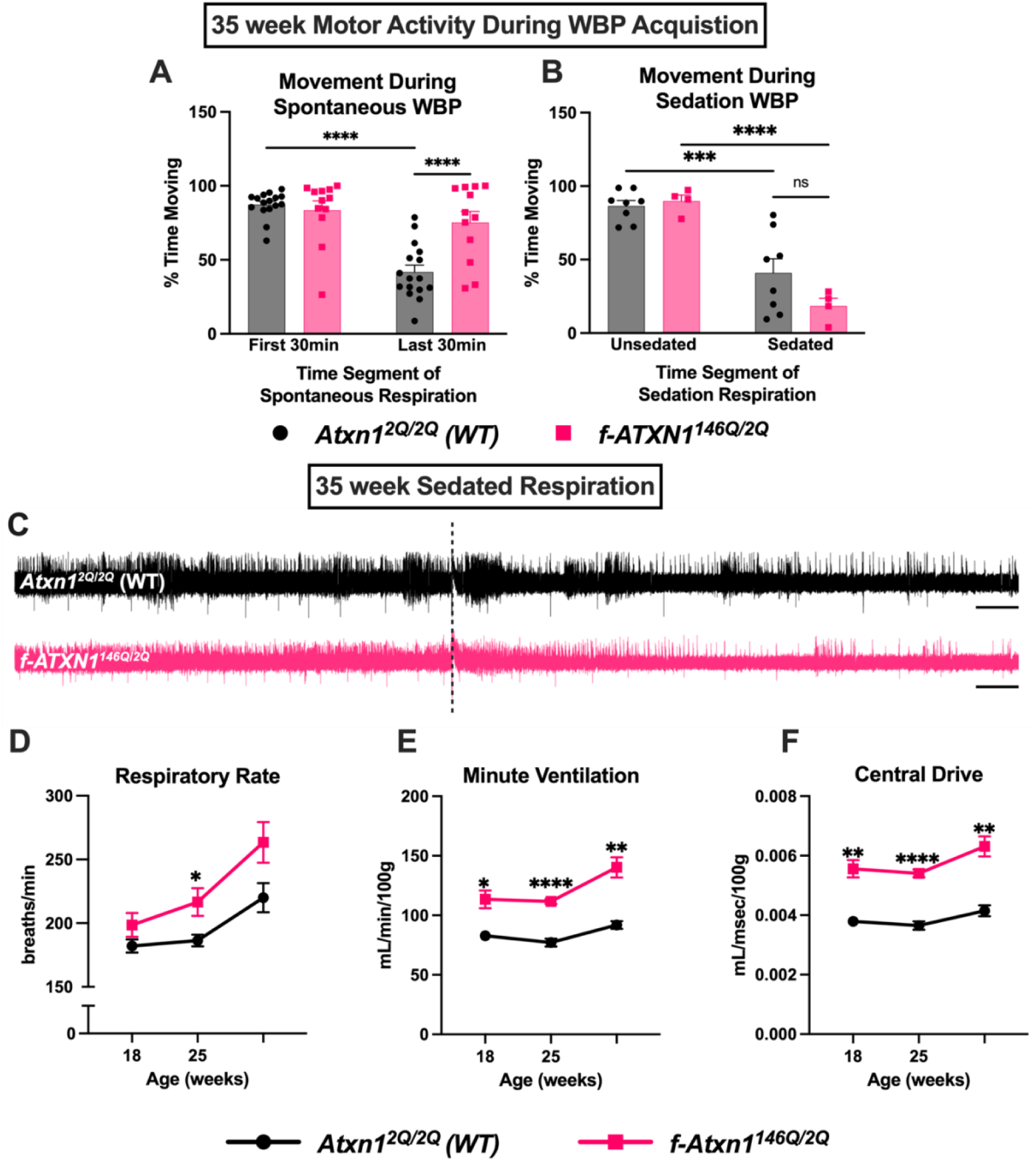
Increased motor activity during WBP is ameliorated with isoflurane sedation indicating functional dysregulation of respiration in *f-ATXN1^146Q/2Q^* mice. (**A**) Movement was quantified during the first and last 30 minutes of spontaneous respiration in WT and *f-ATXN1^146Q/2Q^* mice as the percentage of time spent moving. (**B**) Movement quantified during the first 3 minutes of non-sedated and last 3 minutes of isoflurane sedated respiration in WT and *f-ATXN1^146Q/2Q^* mice. (**C**) WBP flow traces depicting non-sedated respiration followed by the start of isoflurane sedated respiration marked with a dashed line. Scale bar = 1 minute. Sedated respiration was quantified as respiratory rate (**D**), minute ventilation €, and central drive (**F**). Two-way ANOVA for A-B. Mixed effects analysis for D-F with Tukey’s and Geisser-Greenhouse corrections. * p<0.05, ** p<0.01, *** p<0.001, **** p<0.0001. Sample sizes for sedation WBP at each age for WT are 18wk=5, 25wk=8, 35wk=7, and for *f-ATXN1^146Q/2Q^* n=5 for all ages.

To examine respiratory function without the influence of motor behavior, respiration was assessed under a light sedation with isoflurane to reduce exploratory and grooming movements. Isoflurane is a volatile anesthetic that is commonly used in rodent research that has less of an impact on spontaneous respiration compared to other anesthetics (Hao et al., 2021; Massey & Richerson, 2017). WBP was performed under a light sedative dose of isoflurane (0.5%) in normoxia (21% O_2_, 79% N_2_) to examine respiratory function independent of motor activity in WT and *f-ATXN1^146Q/2Q^* mice. Isoflurane sedation was sufficient in reducing movement in WT and *f-ATXN1^146Q/2Q^* mice given both genotypes exhibited reduced movement in the last 3 minutes of sedation compared with the first 3 minutes of non-sedated respiration (Figure 3B). The reduction in movement was associated with a reduction in the amplitude and frequency of respiratory flow traces acquired during sedation (Figure 3C). Respiration under isoflurane respiration reveals a similar, but less severe, phenotype in *f-ATXN1^146Q/2Q^* mice characterized by an increase in the respiratory rate, minute ventilation, and central drive of respiration (Figure 3D-F). Together, this indicates that spontaneous respiratory dysfunction is associated with increased motor activity during the acquisition of WBP in *f-ATXN1^146Q/2Q^* mice. Furthermore, this suggests that there are additional mechanisms beyond behavioral dysfunction that underlie the sedated respiratory phenotype.

### Neural deletion of mutant *ATXN1* in *f-ATXN1^146Q/2Q^* mice ameliorates spontaneous and challenged breathing dysfunction

To assess the extent to which expression of mutant *ATXN1* in respiratory relevant tissues contributes to respiratory dysfunction in *f-ATXN1^146Q/2Q^* mice, these mice were bred to *Nestin-Cre and Acta1-Cre* mice to delete mutant *ATXN1* within neural cells of the brain and spinal cord and skeletal muscle respectively (Duvick et al., 2024). Spontaneous respiration and respiration challenged with hypoxia and hypercapnia were examined using WBP at 25 and 35 weeks as at these ages we observed a significant phenotype in *f-ATXN1^146Q/2Q^* mice. By 35 weeks, *Acta1-Cre;f-ATXN1^146Q/2Q^* mutants exhibited respiratory waveforms with high amplitude and frequency as compared to *f-ATXN1^146Q/2Q^* mice, while the flow traces of *Nestin-Cre;f-ATXN1^146Q/2Q^* mice resembled WT mice (Figure 4A). At 25 and 35 weeks, *Nestin-Cre;f-ATXN1^146Q/2Q^* mice exhibited an amelioration of spontaneous respiratory dysfunction including in the respiratory rate, minute ventilation, and central drive of respiration (Figure 4B-D, Supplemental figure 3C-G). In contrast, *Acta1-Cre;f-ATXN1^146Q/2Q^* mutants exhibited a comparable, and sometimes even exacerbated, spontaneous respiratory dysfunction compared to SCA1 mutants including an elevation in the rate, volume, and central drive of breathing (Figure 4B-D, Supplemental figure 3C-G). The exacerbated respiratory phenotype of *Acta1-Cre;f-ATXN1^146Q/2Q^* mice is associated with an increase in motor activity compared to *f-ATXN1^146Q/2Q^* mice (Supplemental figure 3B). During the respiratory challenge, *Nestin-Cre;f-ATXN1^146Q/2Q^* mice were able to increase respiratory output from spontaneous levels to a similar extent as WT animals (Figure 4E-G, Supplemental figure 3H-J). On the other hand, *Acta1-Cre;f-ATXN1^146Q/2Q^* mice exhibited an elevated respiratory output during spontaneous respiration and failed to further increase respiration when challenged (Figure 4E-G, Supplemental figure 3H-J).

**Figure 4:**
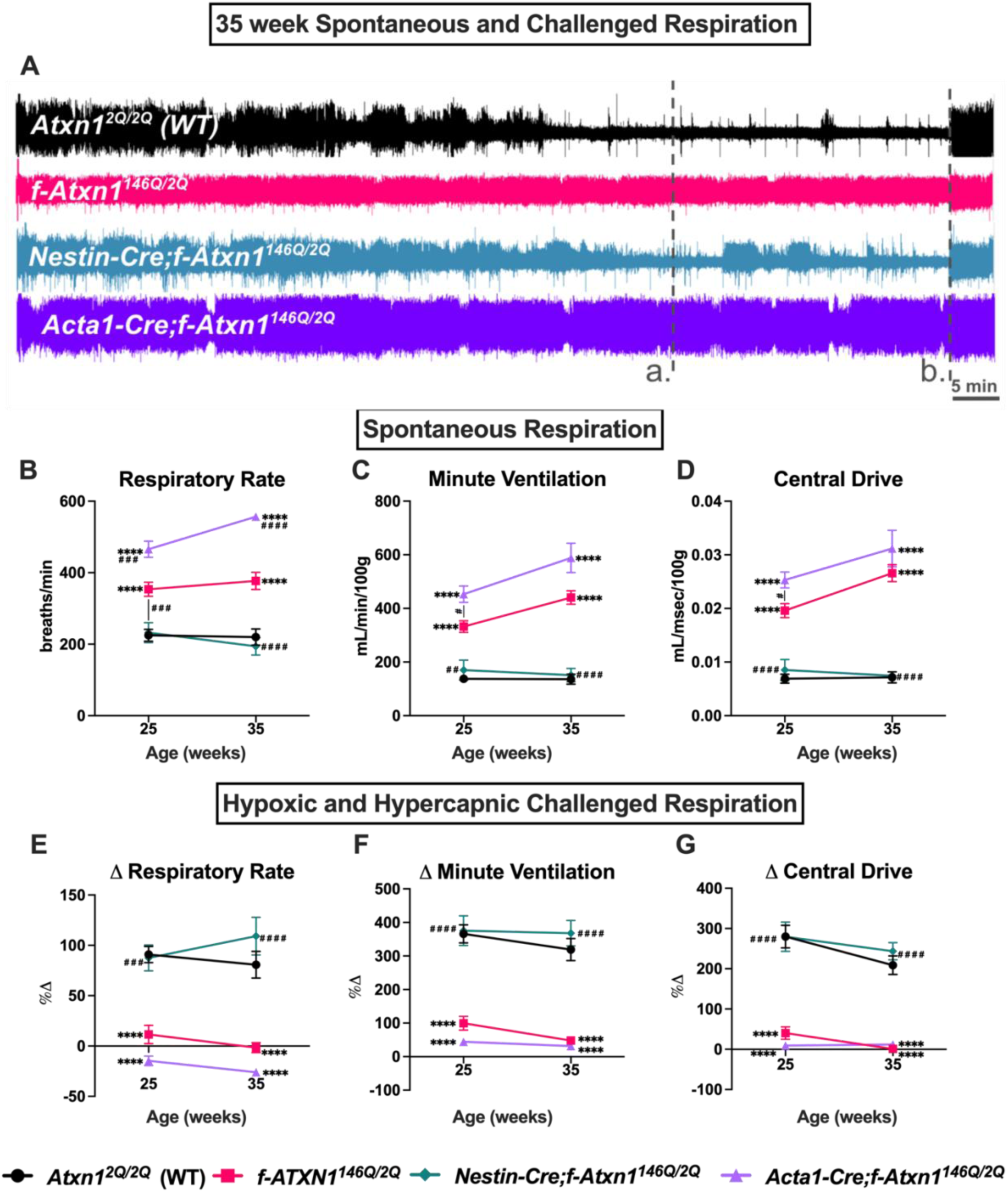
Neural deletion of mutant *ATXN1* in *f-ATXN1^146Q/2Q^* mice ameliorates spontaneous and challenged breathing dysfunction. (**A**) WBP flow traces of spontaneous and challenged respiration. Dashed line a. marks the start of the final 30 minutes of spontaneous respiration while b. marks the beginning of the challenge. Spontaneous respiration was quantified as respiratory rate (**B**), minute ventilation (**C**) and central drive (**D**). The response of mice to a hypoxic and hypercapnic challenge was assessed as the percentage change in respiratory rate €, minute ventilation **(F**) and central drive (**G**) from spontaneous levels. Statistics for B-G were assessed with a repeated measures mixed effects analysis with Tukey’s correction. Asterisks (*) denotes significance at one age compared to WT mice while number signs (#) denote significance at one age compared to *f-ATXN1^146Q/2Q^* mice. */# p<0.05, **/## p<0.01, ***/### p<0.001, ****/#### p<0.0001. WBP sample sizes at each age for WT are 25wk=24, 35wk=21, for *f-ATXN1^146Q/2Q^* are 25wk=19, 35wk=12, for Nestin-Cre:*f-ATXN1^146Q/2Q^* are 25wk=8, 35wk=8, and for Acta1-Cre;*f-ATXN1^146Q/2Q^* mice are 25wk=10, 35wk=6.

Recently, the cerebellum has emerged as a potential player in the regulation of respiration given its connectivity with other respiratory control regions as well as basic and clinical studies that indicate cerebellar pathology can impair respiration (Krohn et al., 2023). As cerebellar dysfunction is a hallmark of SCA1, transgenic *ATXN1[82Q]* mice with Purkinje cell specific expression of mutant ATXN1 were assessed during spontaneous and challenged respiration at 5, 12, and 20 weeks. At all ages assessed, there were no significant differences in respiration observed indicating that Purkinje cells dysfunction is not a major component of respiratory dysfunction caused by mutant *ATXN1* (Supplemental figure 4).

The autonomic regulation of respiration is generated by groups of nuclei in the brainstem, including a subset patterned by the expression of the transcription factor Paired-like homeobox 2b gene (*Phox2b*) during development. Phox2b-Cre mice have enriched expression of Cre in the nucleus of the solitary tract, vagal nucleus, and in the nucleus ambiguus (Scott et al. 2011), regions involved in the chemosensation and motor control of respiration. *F-ATXN1^146Q/2Q^* mice were bred with *Phox2b-Cre* mice to assess the extent that deleting mutant *ATXN1* in Phox2b+ respiratory regions improves respiration. *ATXN1* deletion in *Phox2b-Cre;f-ATXN1^146Q/2Q^* mice was assessed with qPCR for *ATXN1* mRNA extracted from the pons, medulla, and the cervical spinal cord. *Phox2b-Cre;f-ATXN1^146Q/2Q^* mice exhibit a significant reduction of *ATXN1* expression in the cervical spinal cord and trending reductions in the medulla and pons (Supplemental Figure 5B). *Phox2b-Cre;f-ATXN1^146Q/2Q^* mice exhibit a similar respiratory phenotype to *f-ATXN1^146Q/2Q^* mice displaying elevated ventilatory frequency and respiratory output during spontaneous respiration as well as a failure to increase respiratory output during hypoxic and hypercapnic challenge (Supplemental figure 5C-H).

### A neural basis for increased motor activity and sedated respiratory dysfunction in *f-ATXN1^146Q/2Q^* mice that is associated with Phox2b+ respiratory centers

Analysis of the videography acquired during spontaneous respiration reveals that *f-ATXN1^146Q/2Q^* mice exhibit an abnormal behavioral phenotype characterized by an increase in movement during WBP (Figure 3A). To examine the contribution of *ATXN1* within neural tissues to the behavioral dysregulation during WBP, movement was examined in *Nestin-Cre;f-ATXN1^146Q/2Q^* mice in the first and last 30 minutes of videography recorded during spontaneous WBP acquisition. At 35 weeks, *Nestin-Cre;f-ATXN1^146Q/2Q^* mice exhibited similar motor activity as WT and *f-ATXN1^146Q/2Q^* mice in the first 30 minutes of spontaneous respiration. In the final 30 minutes of spontaneous respiration, *Nestin-Cre;f-ATXN1^146Q/2Q^* mice displayed an amelioration of the behavioral phenotype and their movement was reduced to a similar extent as WT mice (Figure 5A). Together this indicates that respiratory dysfunction is associated with increased motor activity during acquisition of spontaneous respiration and that mutant *ATXN1* expression in neural lineages underlies this behavioral dysregulation.

**Figure 5:**
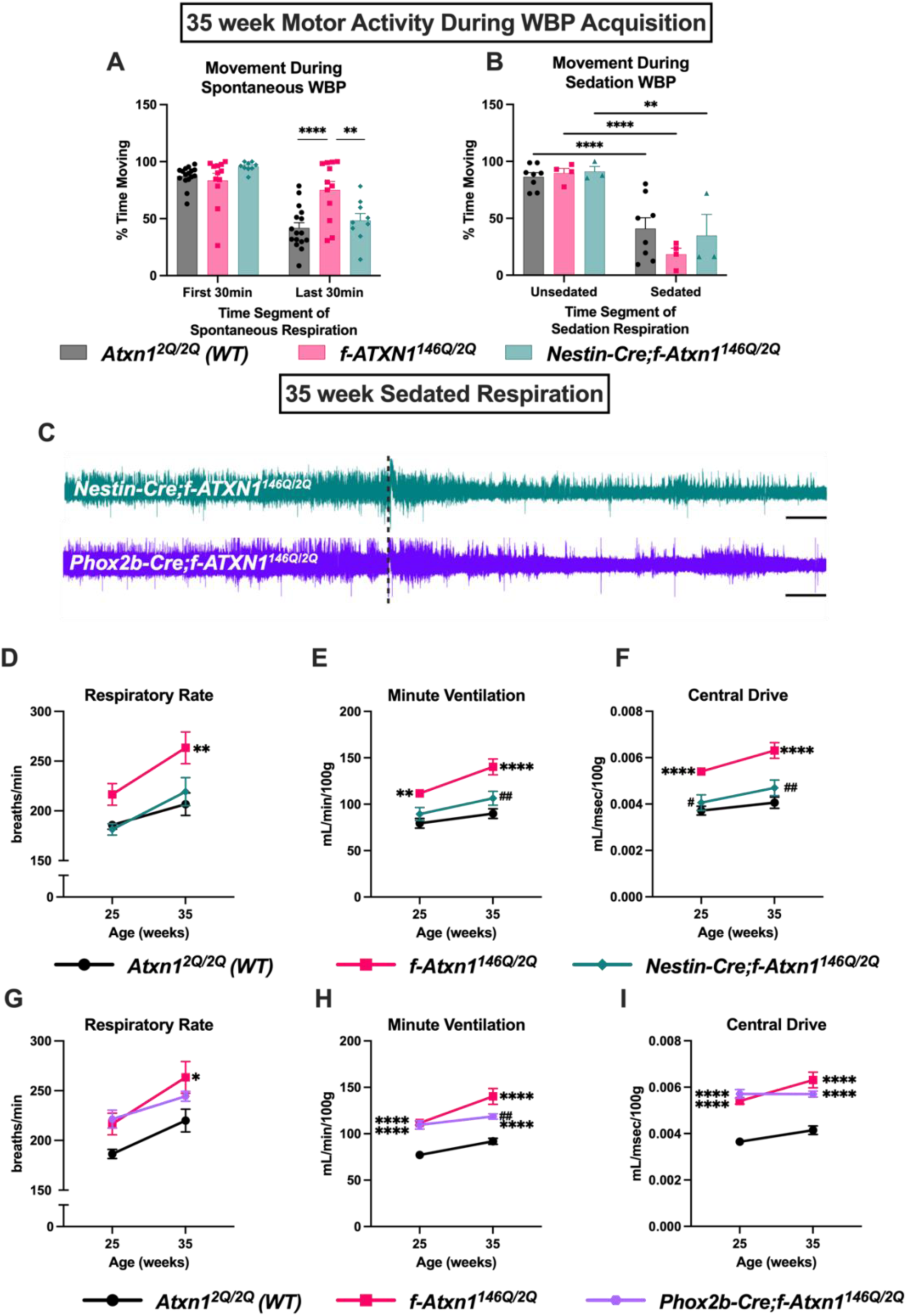
A neural basis for increased motor activity and sedated respiratory dysfunction in *f-ATXN1^146Q/2Q^* mice associated with Phox2b+ respiratory centers. (**A**) Movement was quantified during the first and last 30 minutes of spontaneous respiration as the percentage of time spent moving. (**B**) Movement quantified during the first 3 minutes of non-sedated and last 3 minutes of isoflurane sedated respiration. (**C**) Representative flow traces depicting unsedated respiration followed by isoflurane sedated respiration that is marked with a dashed line. Scale bar = 1 minute. Respiration during sedation with 0.5% isoflurane was quantified as respiratory rate (**D,G**), minute ventilation (**E,H**) and central drive (**F,I**). Repeated measures mixed effects analysis with Tukey’s correction. Asterisks (*) denotes significance at one age compared to WT mice while number signs (#) denote significance at one age compared to *f-ATXN1^146Q/2Q^* mice. */# p<0.05, **/## p<0.01, ***/### p<0.001, ****/#### p<0.0001. Sample sizes for sedation WBP for WT=8, for *f-ATXN1^146Q/2Q^*=5, for *Nestin-Cre:f-ATXN1^146Q/2Q^*=3, and for *Phox2b-Cre:f-ATXN1^146Q/2Q^*=3 at all ages.

To investigate if abnormal sedated respiration is due to a direct effect of mutant *ATXN1* expression in neural lineages, sedated respiration in *Nestin-Cre;f-ATXN1^146Q/2Q^* mice was assessed. Isoflurane sedation was sufficient to reduce motor activity in *Nestin-Cre;f-ATXN1^146Q/2Q^* mice at 35 weeks (Figure 5B). Neural deletion of the mutant *ATXN1* allele attenuated the elevated respiratory phenotype and improved respiratory output at both 25 and 35 weeks demonstrating a large neural contribution to the sedated phenotype (Figure 5D-F).

Given *Phox2b-Cre;f-ATXN1^146Q/2Q^* mice exhibited elevated motor activity during WBP acquisition (Supplemental figure 5A), sedation WBP was employed to examine the role of Phox2b+ cells without the influence of movement. At 25 weeks, *Phox2b-Cre;f-ATXN1^146Q/2Q^* mice exhibited an elevated respiratory rate, minute ventilation, and central drive like *f-ATXN1^146Q/2Q^* mice (Figure 5G-I). *Phox2b-Cre;f-ATXN1^146Q/2Q^* mice exhibited a partial amelioration of minute ventilation at 35 weeks (Figure 5H). Additionally, *Phox2b-Cre;f-ATXN1^146Q/2Q^* mice displayed a trending improvement in central drive compared to *f-ATXN1^146Q/2Q^* mice at 35 weeks (p=0.0868). This suggests that mutant *ATXN1* expression in Phox2b+ regions contributes to abnormal sedated respiration, at least later in disease.

## Discussion

Here, we show the utility of *f-ATXN1^146Q/2Q^* mice to examine regional susceptibility to mutant *ATXN1* and its effect on respiratory dysfunction assessed with whole body plethysmography (WBP). We found that respiratory dysfunction in *f-ATXN1^146Q/2Q^* mice is progressive and age-dependent, becoming especially apparent later in disease. This observation is similar to SCA1 patients that experience motor complications in the initial stages of the disease, and additional symptoms like breathing dysfunction that appear after disease progression (Globas et al., 2008). We demonstrate that spontaneous breathing dysfunction in *f-ATXN1^146Q/2Q^* mice involves multiple mechanisms including a behavioral dysregulation characterized by increased motor activity during acquisition of respiratory function with WBP that is associated with the neural expression of mutant *ATXN1.* Skeletal muscle deletion of mutant *ATXN1* exacerbates the movement and spontaneous respiratory dysfunction phenotypes, and we speculate this is related to improved muscle strength and mobility that is characteristic of this strain (Duvick et al., 2024). A potential neural mechanism for this behavioral phenotype could be related to mood dysregulation that is observed in rodent models of SCA1 (Asher et al., 2021; Tichanek, 2023; Asher et al., 2020) and in SCA patients (Moro et al., 2019; Schmitz-Hübsch et al., 2010, 2011). The ability of isoflurane sedation to ameliorate the behavioral phenotype supports altered mood as a potential mechanism given the depressive effect of sedatives on consciousness. Further research is needed to explore the specific neural mechanisms that underlie this abnormally active behavior observed in mouse models of SCA1 during spontaneous respiration.

Prior to this study, the capability of SCA1 mutant mice to respond to respiratory challenges had not been studied. Mutant *ATXN1* induces a progressive respiratory dysfunction during challenge characterized by a failure to increase ventilation from spontaneous respiration to the same extent as controls. Mutant *ATXN1* may not alter the ability of *f-ATXN1^146Q/2Q^* mice to *sense* changes in oxygen and carbon dioxide levels given that mice were able to increase ventilation during the challenge at earlier ages, but altered chemosensation in aged *f-ATXN1^146Q/2Q^* mice could explain the deficits observed later in disease. The failure of *f-ATXN1^146Q/2Q^* mice to increase respiratory output from spontaneous levels in this study is likely a consequence of the rapid spontaneous respiration in mutant mice without challenge that reduces the dynamic range of respiration and limits additional respiratory capacity. The amelioration of spontaneous respiration and the rescued ability of *Nestin-Cre;f-ATXN1^146Q/2Q^* mice to increase respiration in response to the challenge supports this hypothesis.

Recently, an appreciation for the role of the cerebellum outside of motor coordination has increasingly grown, including in the control of spontaneous and challenged respiration (Fujita et al., 2020; Krohn et al., 2023; Liu et al., 2020; Xu et al., 1995). Given Purkinje cell pathology is a hallmark of SCA1 patients and mouse models, respiration was assessed in the transgenic *ATXN1[82Q]* model of SCA1 with *ATXN1* expression limited to Purkinje cells. Spontaneous and challenged respiration in these mice was unaltered indicating a minor role for Purkinje cells in the abnormal respiratory phenotype of SCA1 mouse models.

Isoflurane sedated WBP allows for respiration to be examined without the influence of motor activity to assess mechanisms of respiratory dysfunction beyond the behavioral phenotype. Given that sedated respiration reveals a similar, but less extreme, respiratory phenotype in *f-ATXN1^146Q/2Q^* mice, we conclude that behavioral dysregulation is a major component of respiratory dysfunction in SCA1 mice and that there are additional mechanisms affecting the respiratory circuitry that contribute to breathing dysfunction. The sedated respiratory phenotype in *f-ATXN1^146Q/2Q^* mice involves inherent dysfunction of central respiratory centers given the substantial amelioration of sedated respiration in *Nestin-Cre;f-ATXN1^146Q/2Q^* mice. Furthermore, respiratory centers expressing Phox2b are a part of the neural amelioration of respiration, at least later in disease. The location of Phox2b+ cell types involved in this rescue are likely located in the cervical spinal cord given the significant reduction of the ATXN1 allele observed in this region. Pons and medulla involvement is also possible given the trending reduction in mutant *ATXN1* observed in these regions.

In summary, we have found evidence that neural expression of mutant *ATXN1* is a major contributor to abnormal respiration in mouse models of SCA1. The partial amelioration of sedated respiration observed by deleting *ATXN1* from neural cells and a subset of cells in the brainstem and spinal cord is consistent with results from recent studies on respiration in SCA1 that associate brainstem and spinal cord degeneration with respiratory dysfunction in rodent models and SCA1 patients (Biswas et al., 2022; Coarelli et al., 2023; Orengo et al., 2018). This research supports the necessity for therapies to reach the brainstem and spinal cord to treat respiratory related complications in SCA1.

## Supplemental Figures

**Supplemental figure 1: *f-ATXN1^146Q/2Q^* extended respiratory parameters.** Spontaneous respiration was quantified as inspiratory time (**A**) and expiratory time (**B**). Tidal volume (**C**), minute ventilation (**D**), and central drive (**E**) are demonstrated without weight adjustment. Respiration during hypoxic challenge was quantified as the percentage change in inspiratory time (**F**), expiratory time (**G**), and tidal volume (**H**) from spontaneous levels. Repeated measures mixed effects analysis with Tukey’s correction for multiple comparisons. * p<0.05, ** p<0.01, *** p<0.001, **** p<0.0001. Sample sizes at each age for WT are 5wk=8, 12wk=6, 18wk=5, 25wk=24, 35wk=21 and for *f-ATXN1^146Q/2Q^* are 5wk=10, 12wk=7, 18wk=5, 25wk=19, 35wk=12.

**Supplemental figure 2: *f-ATXN1^146Q/2Q^* mice exhibit reduced body and lung weights.**

**(A)** Body weight of WT and *f-ATXN1^146Q/2Q^* mice. (**B)** Lung weight from 35-week WT and *f-ATXN1^146Q/2Q^* mice. A repeated measures mixed effects analysis with Geisser Greenhouse and Tukey’s corrections was used for data in A. Student’s t-test for data in B. * p<0.05, *** p<0.001, **** p<0.0001.

**Supplemental figure 3: *Nestin-* and *Acta1-Cre* mediated deletion of mutant *ATXN1* extended respiratory parameters.**

(**A**) Body weights acquired after WBP testing. (**B**) Movement quantified during the first 30 and last 30 minutes of WBP. Spontaneous respiration was quantified as inspiratory time (**C**) and expiratory time (**D**). Tidal volume (**E**), minute ventilation (**F**), and central drive (**G**) are demonstrated without weight adjustment. Respiration during respiratory challenge was quantified as the percentage change in inspiratory time (**H**), expiratory time (**I**), and tidal volume (**J**) from spontaneous levels. Repeated measures mixed effects analysis with Tukey’s correction for multiple comparisons. * p<0.05, ** p<0.01, *** p<0.001, **** p<0.0001. WBP sample sizes at each age for WT are 25wk=24, 35wk=21, for *f-ATXN1^146Q/2Q^* are 25wk=19, 35wk=12, for Nestin-Cre:*f-ATXN1^146Q/2Q^* are 25wk=8, 35wk=8, and for Acta1-Cre;*f-ATXN1^146Q/2Q^* mice are 25wk=10, 35wk=6.

**Supplemental figure 4: Purkinje cell dysfunction does not alter normoxic or challenged respiration.**

Spontaneous respiration in WT and transgenic *PCP2-ATXN1[82Q]* mice was assessed as respiratory rate (**A**), minute ventilation (**B**), and central drive (**C**). Challenged respiration was assessed as percentage change from spontaneous levels for respiratory rate (**A**), minute ventilation (**B**), and central drive (**C**). Repeated measures mixed effects analysis with Geisser Greenhouse and Tukey’s corrections. Sample size for WT at 5wk=9, 12wk=9, and 20wk=6 while for ATXN1[82Q] 5wk=11, 12wk=11, and 20wk=5.

**Supplemental figure 5: Phox2b-Cre mediated ATXN1 deletion does not ameliorate non-sedated respiration.**

(**A**) Movement was quantified during the first and last 30 minutes of spontaneous respiration as the percentage of time spent moving. (**B**) *ATXN1 mRNA* expression in *f-ATXN1^146Q/2Q^* mice and *Phox2b-cre;f-ATXN1^146Q/2Q^* mice. Spontaneous respiration was quantified as respiratory rate (**C**), minute ventilation (**D**), and central drive (**E**). Challenged respiration was assessed as percentage change from spontaneous levels for respiratory rate (**F**), minute ventilation (**G**), and central drive (**H**). Statistics in A are a two-way ANOVA with Tukey’s correction and in B are unpaired t-tests. Statistics in C-E are mixed effects analyses with Geisser Greenhouse and Tukey’s corrections. * p<0.05, ** p<0.01, *** p<0.001, **** p<0.0001. WBP sample sizes at each age for WT are 25wk=24, 35wk=21, for *f-ATXN1^146Q/2Q^* are 25wk=19, 35wk=12, for *Phox2b-cre;f-ATXN1^146Q/2Q^* are 25wk=6, 35wk=8.

## Methods

### Mice

All procedures were approved by the Institutional Animal Care and Use Committee at the University of Minnesota. Mice were housed and managed by Research Animal Resources under specific pathogen free conditions in an AAALAC accredited facility with a 12 hour light/dark cycle and were provided with ad libitum access to food and water. Beginning at 30 weeks all mice within group caging containing *f-ATXN^146Q/2Q^* mice were given DietGel 3M as a supplement to standard dry pellets provided. Animals were age matched within experiments and littermate controls were used when possible. All mouse lines used in this experiment were maintained on a C57B6/J or FVB background and males and females were used in all experiments when possible. WT analyses across all experiments include *Acta1-Cre;Atxn1^2Q/2Q^* and *Nestin-Cre;Atxn1^2Q/2Q^* animals.

### Whole Body Plethysmography

Respiratory function was measured in unrestrained and awake mice with whole body plethysmography (Data Sciences International, St Paul, MN, USA) in a dedicated room separate from shared lab space. Mice were carted to the plethysmography room and allowed to habituate to the space for at least 30 minutes prior to testing. Rectal temperatures were obtained immediately prior to loading mice into clear plexiglass chambers measuring 10cm in diameter. A maximum of six mice were tested in each session, and the view of other testing subjects was obstructed by towels. Spontaneous respiration was first assessed by delivering 21% O_2_ balanced with N_2_ for two hours until a hypoxic and hypercapnic challenge consisting of 5% CO_2_, 10% O_2_, and a balance of N_2_ was automatically initiated for a total duration of 5 minutes. At the end of the recording, rectal temperatures and body weights were recorded and mice were placed back in their home cages.

WBP analysis was performed with Ponemah software (Data Sciences International, St. Paul, MN, USA). To quantify spontaneous respiration during normoxia, a 3-minute segment of respiration was selected from the final 30 minutes of normoxia that represented the most frequent and consistent respiratory pattern in that timeframe. This region was chosen given that WT mice took up to 1.5 hours to exhibit quiet resting, defined as periods of behavior with absent exploratory or grooming behaviors. To quantify the respiratory challenge the final minute of the challenge was analyzed across all individuals. Ventilatory parameters including respiratory rate, inspiratory and expiratory times, tidal volume and minute ventilation were automatically calculated from parsed data. Central drive was calculated as the quotient of tidal volume and inspiratory time and expressed as mL/g/sec^-1^. Tidal volumes, minute ventilation, and central drive were all normalized to body weight.

### Whole Body Plethysmography with Isoflurane Sedation

Whole body plethysmography (Data Sciences International, St Paul, MN, USA) was used to assess ventilation in unrestrained mice lightly sedated with isoflurane. Rectal temperatures were obtained immediately prior to loading the mice into one of the two plethysmography chambers used for this experiment. Initially mice were acclimated to the chamber in normoxic conditions (21% O_2_ balanced with N_2_) for 10 minutes which was followed by a 10-15-minute recording with 0.5% isoflurane mixed in 21% O_2_ and a balance of N_2_. With this protocol mice took on average 5 minutes to exhibit sedated behavior after the induction of isoflurane that was characterized by a quiet resting behavior absent of grooming and exploration. After the conclusion of isoflurane administration, mice were allowed to recover in normoxia for 10 minutes before they were removed from the chamber and their rectal temperatures and body weights were recorded.

WBP analysis was performed with Ponemah software (Data Sciences International, St. Paul, MN, USA). Data selected for automatic analysis included a 3 minute segment of respiration that represented sedated quiet respiration with minimal movement. Mice exhibited spontaneous jerky movements under light sedation, so segments of respiration with movement or sniffing sequences were removed from the quantification.

### Videography Movement Analysis

Videography acquired during spontaneous plethysmography acquisition was analyzed for immobility duration using ANY-maze video tracking software (Stoeling Co. Wood Dale, IL, USA). Immobility was assessed in the first 30 minutes and final 30 minutes of the two hour spontaneous respiration recording. The minimum duration for immobility episodes to be counted was 4 seconds. To quantify the duration of time moving, the duration of time immobile for each mouse was subtracted from the total 30 minutes and the percentage of time moving was calculated as the movement duration divided by total time multiplied by 100.

Videography acquired during sedation plethysmography acquisition was analyzed for immobility duration in the first 3 minutes of the non-sedated respiratory recordings and the last 3 minutes of respiration sedated with 0.5% isoflurane. The minimum duration for counted immobility episodes was 2 seconds. To quantify the duration of time moving, the duration of time immobile for each mouse was subtracted from 3 minutes and the percentage of time moving was calculated as the movement duration divided by total time multiplied by 100.

### qPCR

To assess the relative expression of human *ATXN1 mRNA,* qPCR was used as previously described in Duvick et al. 2024. Briefly, microdissected brain regions stored in RNAlater were hydrated with RNase free water and were homogenized with TRIzol (Thermo Fisher Scientific) to isolate RNA. cDNA was synthesized in duplicate using iScript Advanced cDNA synthesis Kit (Bio-Rad) and diluted 1:5 with water. Custom primers for human *ATXN1* were then used to assess *ATXN1* expression relative to the expression of the mouse reference gene GAPDH with the ΔΔCt method of quantification.

### Lung weights

WT and *f-ATXN1^146Q/2Q^* mice aged between 30 and 40 weeks were euthanized by transcardial perfusions and fixed with 10% formalin before the left and right lungs were removed from the abdomen and teased away from any tracheal or fat tissue attached. Extracted lungs were rinsed in PBS, dabbed on filter paper, and then weighed on a balance.

### Statistical Analysis

All statistical analyses were performed using Graphad prism (v10.0). Data in all experiments was assessed to meet the assumptions of the appropriate statistical analysis. All data is represented as 土 the mean standard error of the mean (SEM). In all cases, P < 0.05 was considered statistically significant.

## Acknowledgments

This study was supported by a National Ataxia Foundation grant and NIH grants 1036654 (AS), R01NS109077 (MC), and R35NS127248 (HO) respectively. The authors thank Rebecca Barok for the initial design of plethysmography protocols.

## Author Contributions

A.S., H.O., and M.C. conceived the study, designed experiments, and wrote the paper. A.S., H.O., M.C., B.D., and J.G. interpreted data. A.S. and P.Y. performed acquisition of animal behaviors. A.S., K.D., R.B, and R.C analyzed plethysmography data. K.D., P.Y. and A.S. performed tissue dissections. L.D. extracted RNA and performed qPCR. C.C. performed analyses of movement during plethysmography. S.S. and O.R. bred mice and managed the colony. A.S. performed statistical analyses. All authors reviewed the manuscript.

## References

1. Asher, M., Rosa, J.-G., & Cvetanovic, M. (2021). Mood alterations in mouse models of Spinocerebellar Ataxia type 1. Scientific Reports, 11(1), 713. 10.1038/s41598-020-80664-9

2. Asher, M., Rosa, J.-G., Rainwater, O., Duvick, L., Bennyworth, M., Lai, R.-Y., CRC-SCA, Kuo, S.-H., & Cvetanovic, M. (2020). Cerebellar contribution to the cognitive alterations in SCA1: Evidence from mouse models. Human Molecular Genetics, 29(1), Article 1. 10.1093/hmg/ddz265

3. Biswas, D. D., El Haddad, L., Sethi, R., Huston, M. L., Lai, E., Abdelbarr, M. M., Mhandire, D. Z., & ElMallah, M. K. (2022). Neuro-respiratory pathology in spinocerebellar ataxia. Journal of the Neurological Sciences, 443, 120493. 10.1016/j.jns.2022.120493

4. Bloem, B. R., Okun, M. S., & Klein, C. (2021). Parkinson’s disease. The Lancet, 397(10291), 2284–2303. 10.1016/S0140-6736(21)00218-X

5. Coarelli, G., Tchikviladzé, M., Dodet, P., Arnulf, I., Charles, P., Tankeré, F., Similowski, T., Seilhean, D., Brice, A., Duyckaerts, C., & Durr, A. (2023). Motor neuron involvement threatens survival in spinocerebellar ataxia type 1. Neuropathology and Applied Neurobiology, 49(2), e12897. 10.1111/nan.12897

6. Diallo, A., Jacobi, H., Cook, A., Labrum, R., Durr, A., Brice, A., Charles, P., Marelli, C., Mariotti, C., Nanetti, L., Panzeri, M., Rakowicz, M., Sobanska, A., Sulek, A., Schmitz-Hübsch, T., Schöls, L., Hengel, H., Melegh, B., Filla, A., … Tezenas du Montcel, S. (2018). Survival in patients with spinocerebellar ataxia types 1, 2, 3, and 6 (EUROSCA): A longitudinal cohort study. The Lancet. Neurology, 17(4), 327–334. 10.1016/S1474-4422(18)30042-5

7. Duvick, L., Southern, W. M., Benzow, K. A., Burch, Z. N., Handler, H. P., Mitchell, J. S., Kuivinen, H., Gadiparthi, U., Yang, P., Soles, A., Sheeler, C. A., Rainwater, O., Serres, S., Lind, E. B., Nichols-Meade, T., You, Y., O’Callaghan, B., Zoghbi, H. Y., Cvetanovic, M., … Orr, H. T. (2024). Mapping SCA1 regional vulnerabilities reveals neural and skeletal muscle contributions to disease. JCI Insight, 9(9). 10.1172/jci.insight.176057

8. Fujita, H., Kodama, T., & du Lac, S. (2020). Modular output circuits of the fastigial nucleus for diverse motor and nonmotor functions of the cerebellar vermis. eLife, 9, e58613. 10.7554/eLife.58613

9. Fusco, A. F., Pucci, L. A., Switonski, P. M., Biswas, D. D., McCall, A. L., Kahn, A. F., Dhindsa, J. S., Strickland, L. M., La Spada, A. R., & ElMallah, M. K. (2021). Respiratory dysfunction in a mouse model of spinocerebellar ataxia type 7. Disease Models & Mechanisms, 14(7), dmm048893. 10.1242/dmm.048893

10. Gaig, C., & Iranzo, A. (2012). Sleep-disordered breathing in neurodegenerative diseases. Current Neurology and Neuroscience Reports, 12(2), 205–217. 10.1007/s11910-011-0248-1

11. Genis, D., Matilla, T., Volpini, V., Rosell, J., Davalos, A., Ferrer, I., Molins, A., & Estivill, X. (1995). Clinical, neuropathologic, and genetic studies of a large spinocerebellar ataxia type 1 (SCA1) kindred: (CAG) _n_ expansion and early premonitory signs and symptoms. Neurology, 45(1), Article 1. 10.1212/WNL.45.1.24

12. Georgopoulos, D., Bolaki, M., Stamatopoulou, V., & Akoumianaki, E. (2024). Respiratory drive: A journey from health to disease. Journal of Intensive Care, 12(1), 15. 10.1186/s40560-024-00731-5

13. Globas, C., du Montcel, S. T., Baliko, L., Boesch, S., Depondt, C., DiDonato, S., Durr, A., Filla, A., Klockgether, T., Mariotti, C., Melegh, B., Rakowicz, M., Ribai, P., Rola, R., Schmitz-Hubsch, T., Szymanski, S., Timmann, D., Van de Warrenburg, B. P., Bauer, P., & Schols, L. (2008). Early symptoms in spinocerebellar ataxia type 1, 2, 3, and 6. Movement Disorders: Official Journal of the Movement Disorder Society, 23(15), 2232–2238. 10.1002/mds.22288

14. Hao, X., Ou, M., Li, Y., & Zhou, C. (2021). Volatile anesthetics maintain tidal volume and minute ventilation to a greater degree than propofol under spontaneous respiration. BMC Anesthesiology, 21(1), 238. 10.1186/s12871-021-01438-y

15. Hobson, P., & Meara, J. (2018). Mortality and quality of death certification in a cohort of patients with Parkinson’s disease and matched controls in North Wales, UK at 18 years: A community-based cohort study. BMJ Open, 8(2), e018969. 10.1136/bmjopen-2017-018969

16. Huebra, L., Coelho, F. M., Filho, F. M. R., Barsottini, O. G., & Pedroso, J. L. (2019). Sleep Disorders in Hereditary Ataxias. Current Neurology and Neuroscience Reports, 19(8), Article 8. 10.1007/s11910-019-0968-1

17. Krohn, F., Novello, M., van der Giessen, R. S., De Zeeuw, C. I., Pel, J. J., & Bosman, L. W. (2023). The integrated brain network that controls respiration. eLife, 12, e83654. 10.7554/eLife.83654

18. Larson, T. C., Goutman, S. A., Davis, B., Bove, F. J., Thakur, N., & Mehta, P. (2023). Causes of death among United States decedents with ALS: An eye toward delaying mortality. Annals of Clinical and Translational Neurology, 10(5), 757–764. 10.1002/acn3.51762

19. Liu, Y., Qi, S., Thomas, F., Correia, B. L., Taylor, A. P., Sillitoe, R. V., & Heck, D. H. (2020). Loss of cerebellar function selectively affects intrinsic rhythmicity of eupneic breathing. Biology Open, 9(4), Article 4. 10.1242/bio.048785

20. Massey, C. A., & Richerson, G. B. (2017). Isoflurane, ketamine-xylazine, and urethane markedly alter breathing even at subtherapeutic doses. Journal of Neurophysiology, 118(4), 2389–2401. 10.1152/jn.00350.2017

21. Morgan, B. J., Adrian, R., Bates, M. L., Dopp, J. M., & Dempsey, J. A. (2014). Quantifying hypoxia-induced chemoreceptor sensitivity in the awake rodent. Journal of Applied Physiology, 117(7), 816–824. 10.1152/japplphysiol.00484.2014

22. Moro, A., Moscovich, M., Farah, M., Camargo, C. H. F., Teive, H. A. G., & Munhoz, R. P. (2019). Nonmotor symptoms in spinocerebellar ataxias (SCAs). Cerebellum & Ataxias, 6(1), 12. 10.1186/s40673-019-0106-5

23. Orengo, J. P., van der Heijden, M. E., Hao, S., Tang, J., Orr, H. T., & Zoghbi, H. Y. (2018). Motor neuron degeneration correlates with respiratory dysfunction in SCA1. Disease Models & Mechanisms, 11(2), Article 2. 10.1242/dmm.032623

24. Robitaille, Y., Schut, L., & Kish, S. J. (1995). Structural and immunocytochemical features of olivopontocerebellar atrophy caused by the spinocerebellar ataxia type 1 (SCA-1) mutation define a unique phenotype. Acta Neuropathologica, 90(6), Article 6. 10.1007/BF00318569

25. Sasaki, H., Fukazawa, T., Yanagihara, T., Hamada, T., Shima, K., Matsumoto, A., Hashimoto, K., Ito, N., Wakisaka, A., & Tashiro, K. (1996). Clinical features and natural history of spinocerebellar ataxia type 1. Acta Neurologica Scandinavica, 93(1), 64–71. 10.1111/j.1600-0404.1996.tb00173.x

26. Schmitz-Hübsch, T., Coudert, M., Giunti, P., Globas, C., Baliko, L., Fancellu, R., Mariotti, C., Filla, A., Rakowicz, M., Charles, P., Ribai, P., Szymanski, S., Infante, J., van de Warrenburg, B. P. C., Dürr, A., Timmann, D., Boesch, S., Rola, R., Depondt, C., … Klockgether, T. (2010). Self-rated health status in spinocerebellar ataxia—Results from a European multicenter study. Movement Disorders, 25(5), 587–595. 10.1002/mds.22740

27. Schmitz-Hübsch, T., Coudert, M., Tezenas du Montcel, S., Giunti, P., Labrum, R., Dürr, A., Ribai, P., Charles, P., Linnemann, C., Schöls, L., Rakowicz, M., Rola, R., Zdzienicka, E., Fancellu, R., Mariotti, C., Baliko, L., Melegh, B., Filla, A., Salvatore, E., … Klockgether, T. (2011). Depression comorbidity in spinocerebellar ataxia. Movement Disorders: Official Journal of the Movement Disorder Society, 26(5), 870–876. 10.1002/mds.23698

28. Sørensen, S. A., & Fenger, K. (1992). Causes of death in patients with Huntington’s disease and in unaffected first degree relatives. Journal of Medical Genetics, 29(12), 911–914. 10.1136/jmg.29.12.911

29. Tichanek, F. (2023). Psychiatric-Like Impairments in Mouse Models of Spinocerebellar Ataxias. *Cerebellum (London*, England), 22(1), 14–25. 10.1007/s12311-022-01367-7

30. Xu, F., Owen, J., & Frazier, D. T. (1995). Hypoxic respiratory responses attenuated by ablation of the cerebellum or fastigial nuclei. Journal of Applied Physiology (Bethesda, Md.: 1985), 79(4), Article 4. 10.1152/jappl.1995.79.4.1181

