## Supplemental figures for "A Neural Basis for Mutant ATAXIN-1 Induced Respiratory Dysfunction in Mouse Models of Spinocerebellar Ataxia Type 1"

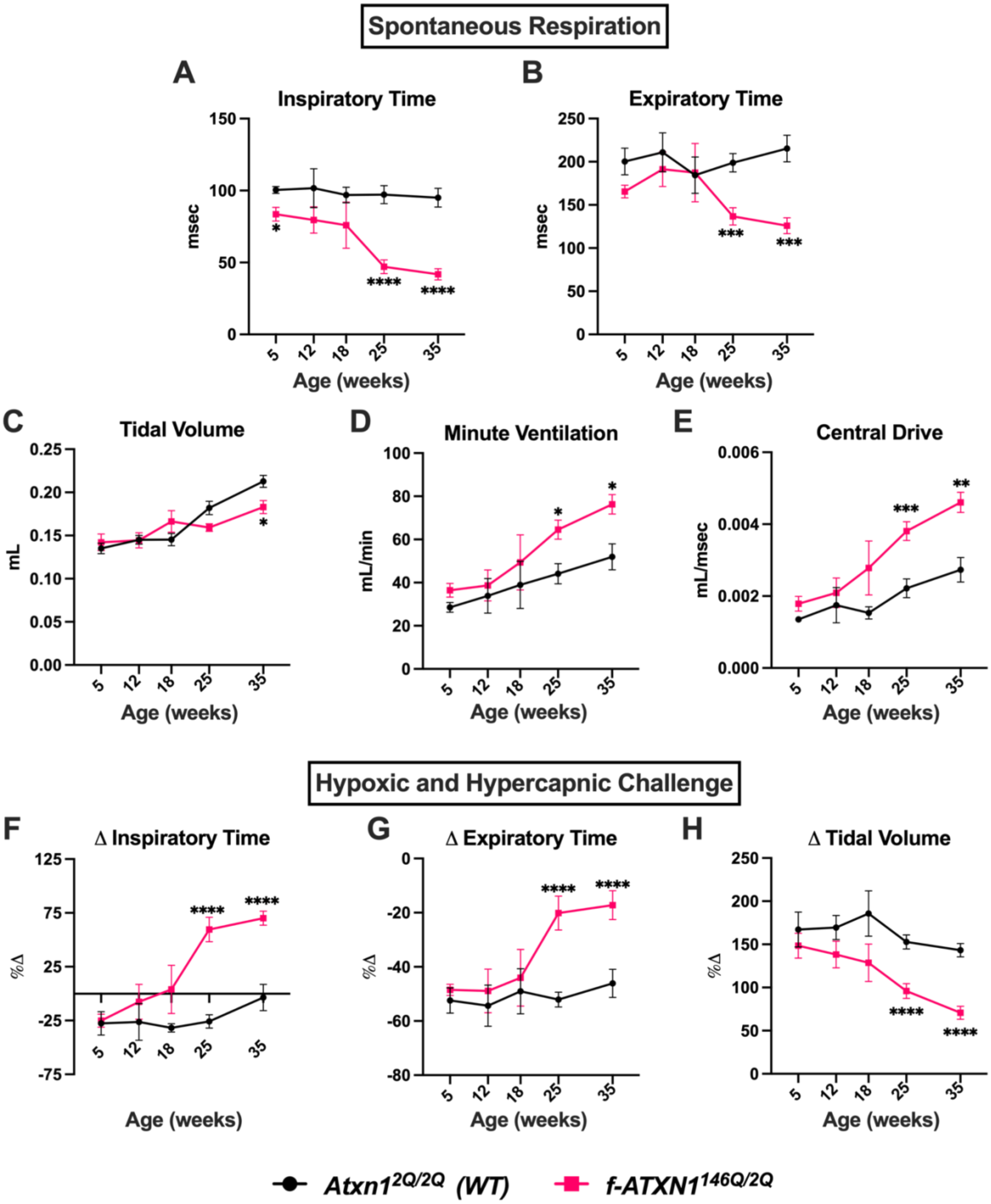


**Supplemental figure 1**

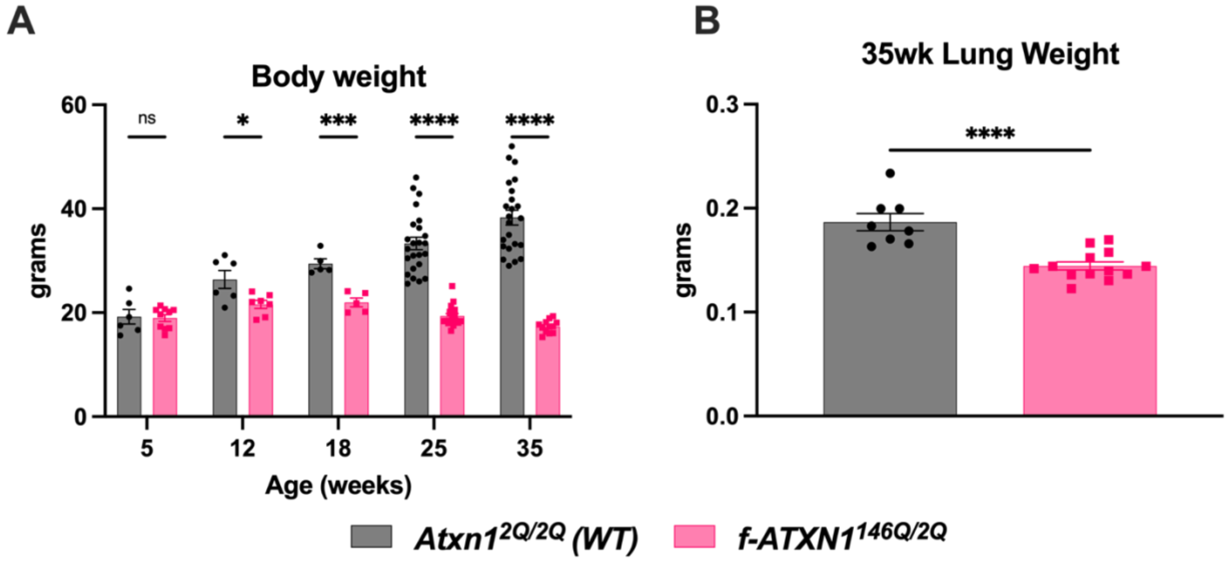


**Supplemental figure 2**

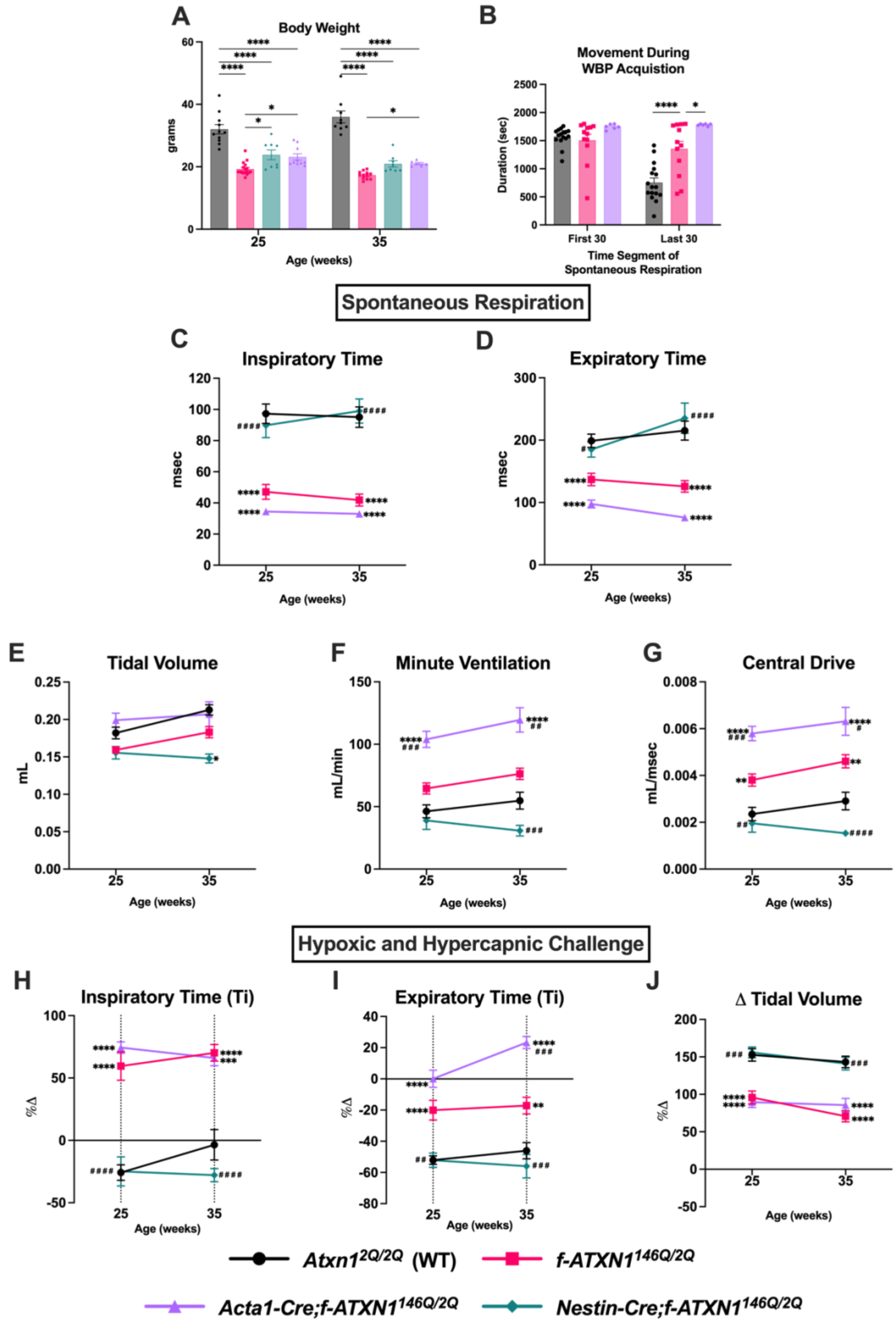


**Supplemental figure 3**

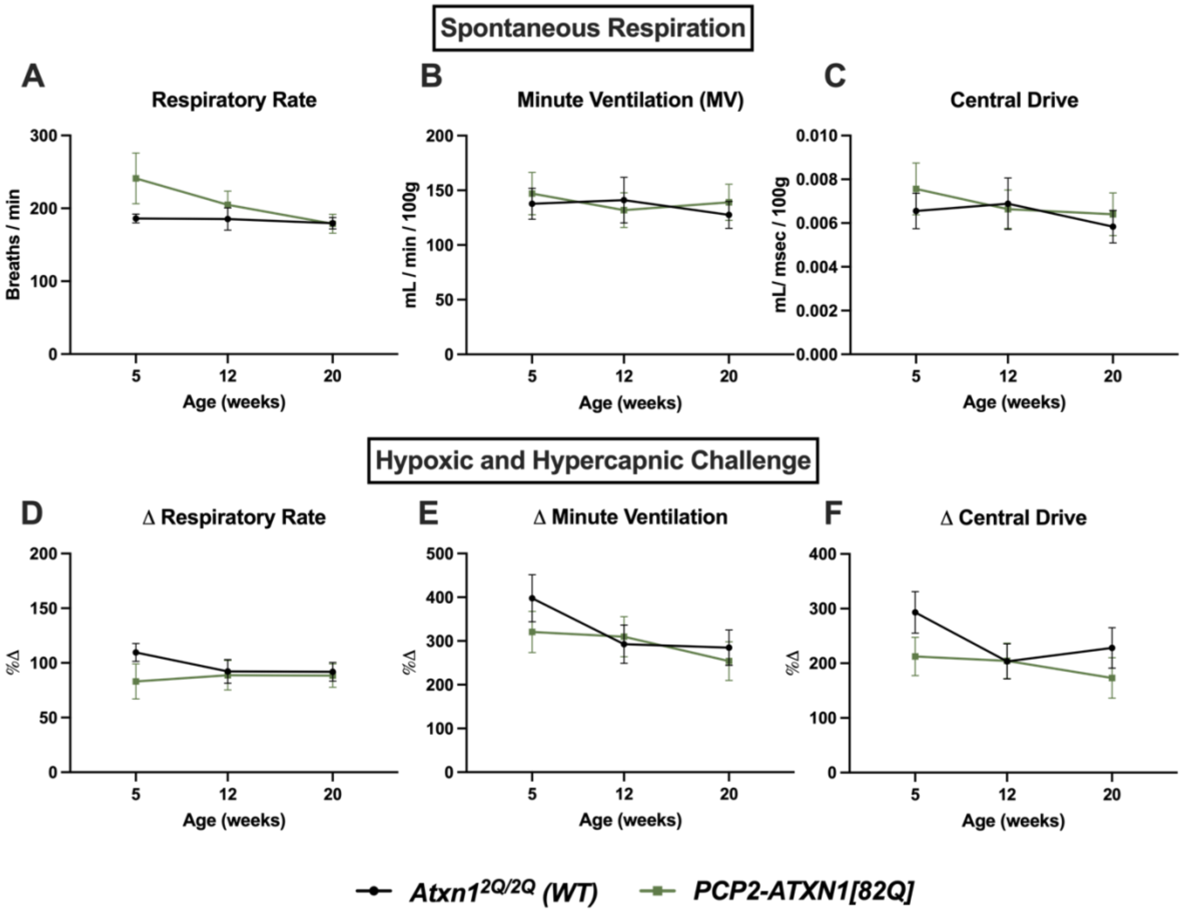


**Supplemental figure 4**

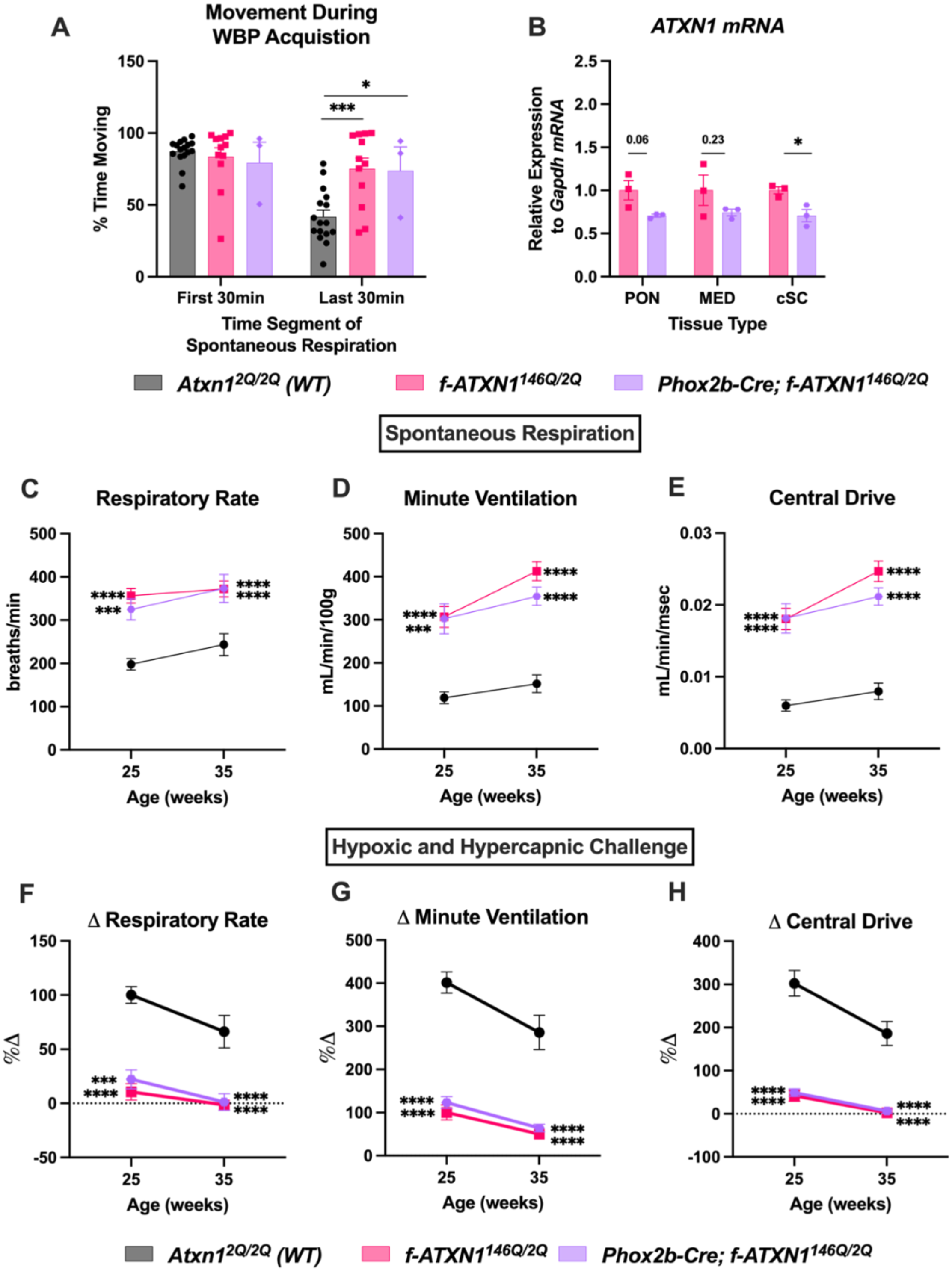


**Supplemental figure 5**
